# S3R: Modeling spatially varying associations with Spatially Smooth Sparse Regression

**DOI:** 10.1101/2025.09.06.674629

**Authors:** Xinyu Zhou, Pengtao Dang, Xiao Wang, Laura Xianlu Peng, Jen Jen Yeh, Nan Zhang, Brian Neelon, Rosie Sears, Teresa Zimmers, Chi Zhang, Sha Cao

## Abstract

Spatial transcriptomics (ST) data demands models that recover how associations among molecular and cellular features change across tissue while contending with noise, collinearity, cell mixing, and thousands of predictors. We present Spatially Smooth Sparse Regression (S3R), a general statistical framework that estimates location-specific coefficients linking a response feature to high-dimensional spatial predictors. S3R unites structured sparsity with a minimum-spanning-tree–guided smoothness penalty, yielding coefficient fields that are coherent within neighborhoods yet permit sharp boundaries. In synthetic data, S3R accurately recovers spatially varying effects, selects relevant predictors, and preserves known boundaries. Applied to Visium-based ST data, S3R recapitulates layer-specific target–TF associations in human dorsolateral prefrontal cortex with concordant layer-wise correlations in matched single-cell data. In acute *Haemophilus ducreyi* skin infection, S3R converts spot-level gene expression mixtures into cell type–attributed expression fields, revealing per-cell type spatial gradients, and improving concordance of spatially variable gene calls when tests are applied to these demixed fields. In pancreatic ductal adenocarcinoma, S3R builds cross–cell-type, cross-gene co-variation tensors that quantify cell–cell interaction strength at gene-pair resolution and nominate interacting genes whose pathway enrichments align with established stromal–epithelial and immune crosstalk. An efficient implementation scales to large assays, and on a Xenium-based breast cancer dataset, S3R delineates the contributions at gene–gene, local neighborhood, and global context-level to target gene expression. Because responses and predictors in S3R are user-defined, it could flexibly address diverse biological questions within a single, scalable, and interpretable regression framework.

## Introduction

Spatial transcriptomics (ST) has transformed our ability to map gene expression in intact tissues, by revealing how cellular neighborhoods and tissue architecture shape molecular phenotypes [1]. By preserving spatial context that is lost in dissociated single-cell assays, ST enables questions that were previously out of reach [2]: how adjacent cell populations communicate [3–6], how a cell’s position within a layer or niche modulates its expression program [7, 8], and how proximity among distinct cell types influences function [9, 10]. Existing computational approaches have delivered important advances, by identifying spatially variable genes and segmenting tissues into expression domains or biological niches [4, 11, 12], yet they often analyze a small set of features at a time, treat locations as discrete blocks, or rely on dimension reduction that can mask important signals [13, 14]. What remains comparatively underexplored is not merely where spatial features are patterned, but how ***relationships*** among spatial features, such as genes, transcriptional regulators, cell-type abundances, local/global microenvironmental contexts, and other covariates, change with location [15, 16].

Addressing this gap in spatial transcriptomics calls for models that estimate location-specific associations while remaining stable enough to be interpretable in the presence of noise, collinearity, and spatial autocorrelation, and respecting sharp boundaries where microenvironments change. These associations are most naturally captured as the coefficients of a multivariable model. For estimating associations, classical regression models treats coefficients as global, assuming stationarity; mixture models allow a few discrete clusters but do not exploit coordinates and often miss the gradual transitions that characterize tissue architecture [17]. High dimensionality further complicates the computation when thousands of spatial features (such as genes) are candidates to explain spatial variation. Although recent high-dimensional spatial regression methods introduce sparsity to perform variable selection [18], they typically assume a single, tissue-wide set of selected predictors and therefore omit spatially varying selection. In practice, the relevant features change with context: regulators of hypoxia may dominate near poorly perfused regions [19]; adhesion and ECM programs may become predictive adjacent to ducts or vasculature [20]; immune signaling may emerge only within specific niches [21].

Consequently, the set of active predictors should be allowed to differ across locations, and their effects should vary smoothly within neighborhoods yet change abruptly at histological interfaces when biology dictates. On the technology side, spot-level measurements aggregate contributions from mixtures of cells in the case of non-single cell resolution ST technologies such as Visium and NanoString GeoMx DSP, so models must mitigate cell-mixing confounding to avoid diluting true gradients and spurious associations. A suitable framework must therefore (i) enforce parsimony for interpretability and stability in high dimensions, (ii) estimate spatially adaptive coefficients and enable location-dependent variable inclusion, (iii) preserve sharp boundaries while borrowing strength within neighborhoods, and (iv) scale computationally to modern ST datasets spanning tens of thousands of features and spatial locations.

Here we introduce Spatially Smooth Sparse Regression (S3R), a general approach for modeling spatially varying associations in high-dimensional ST data. S3R implements structured sparsity, to allow only a small subset of predictors to be retained for interpretability and statistical stability, with spatially adaptive smoothness, that borrows strength within neighborhoods yet permits sharp transitions at histological boundaries. Specifically, S3R estimates location-specific coefficients for each predictor–response pair, and produces spatial fields whose magnitudes encode the strength of association and whose gradients capture continuous trends or abrupt shifts between microenvironmental niches. In addition, S3R performs location-specific variable selection, where the set of active predictors is allowed to change across space, enabling different features to enter (or drop out of) the model in distinct tissue regions while maintaining coherence within each neighborhood. Because responses and predictors can be flexibly defined, such as gene expression as a function of transcription factor expression, cell-type proportions, local microenvironmental composition, tissue-level niche context, etc, S3R could unify several analysis goals within a single, interpretable regression framework.

We applied S3R to two broad classes of tissues: those with relatively well-defined architecture and cellular neighborhoods such as cortex and inflamed skin, and those with highly intermixed cell populations, such as pancreatic and breast cancer tissues. We demonstrate the flexible application of S3R across distinct settings that highlight its strengths. In human dorsolateral prefrontal cortex (DLPFC), where laminar structure is well annotated, S3R recovers spatially coherent, layer-specific associations between targets and transcriptional regulators. In acute *Haemophilus ducreyi* skin infection, S3R converts spot-level mixtures into cell type–attributed expression fields and reveals tissue-scale gradients that are not visible to other methods. In pancreatic ductal adenocarcinoma (PDAC), S3R enables transcriptome-wide, cross–cell type, cross-gene co-variation analysis, and summarizes replicated gene-gene couplings between interacting cell type pairs. Finally, on a Xenium-based breast cancer dataset, S3R scales to single-cell/in situ resolution and delineates gene–gene regulatory patterns together with local-neighborhood and niche-level contributions to expression, illustrating applicability to large images with dense sampling. Across diverse tissues and study designs, S3R offers a unified way to interrogate *how* spatial context shapes biology rather than merely *where* expression is high or low. By estimating location-specific coefficients and allowing the active predictor set to vary across space, it attributes gene programs to specific cell types in situ, dissects transcriptional regulation at the level of transcription factors and at local-and global-context scales, delineates gradients and sharp boundaries of activity, and quantifies gene-pair–anchored co-variation between cell types. The result is a compact, interpretable atlas of spatially varying associations that yields testable hypotheses about cell type–specific regulation, coordinated intercellular crosstalk, and context-defined organization of tissue function.

## Results

### Methods overview

Spatially smooth and Sparse Regression (S3R) is a computational framework designed to resolve various spatially varying relationships in noisy, high-dimensional ST data (**Fig. 1**). The method addresses two critical challenges in modeling the relationship among spatial features: biologically meaningful relationships may not be static across tissue regions, and the spatial features that encode such relationships are frequently masked by measurement noise, collinearity, and a high dimensional predictor set. S3R unifies sparsity-inducing regularization with spatially aware smoothing constraints. At its core, S3R models the relationship between one spatial feature, namely the response feature (**Fig. 1a**), and a set of other spatial features (**Fig. 1b**), namely the predictor features, through spatially adaptive coefficients, 𝜷_𝒌_(𝒔_𝒊_) (**Fig. 1c**), which quantify the influence of each predictor feature 𝒌 at location 𝒔_𝒊_onto the response feature. To ensure robustness, the framework imposes three complementary penalties: 1) a *minimum spanning tree (MST)-based spatial smoothness penalty* on edgewise coefficient differences, that enforces local homogeneity in coefficients across adjacent tissue regions while permitting sharp boundaries (**Fig. 1d**)[22]; 2) an 𝑳_𝟏_ *penalty* on coefficients to induce location-specific sparsity; and 3) a *group* 𝑳_𝟐_ penalty to eliminate non-predictive features globally (**Fig. 1e**). The MST, constructed from spatial coordinates using Euclidean distances, reduces computational complexity by representing tissue topology with only 𝒏 − 𝟏 edges while maintaining full connectivity. Optimization is achieved through the highly efficient Adam optimizer [23] using a scalable multi-GPU parallel framework [24], with hyperparameters tuned in parallel using Optuna [25]. These optimizations enable efficient training and model selection on large-scale ST datasets. On comprehensively configured simulation settings, S3R most faithfully recovered the true location-specific coefficients in terms of the clustering structure, both by visual inspection (Extended Fig. 1a) and by Adjusted Rand Index (ARI; Extended Fig. 1b). This is true for both high dimensional and relatively lower dimensional predictor settings, and different configurations of the spatial heterogeneity. See Methods and Materials for more details.

**Fig. 1.**
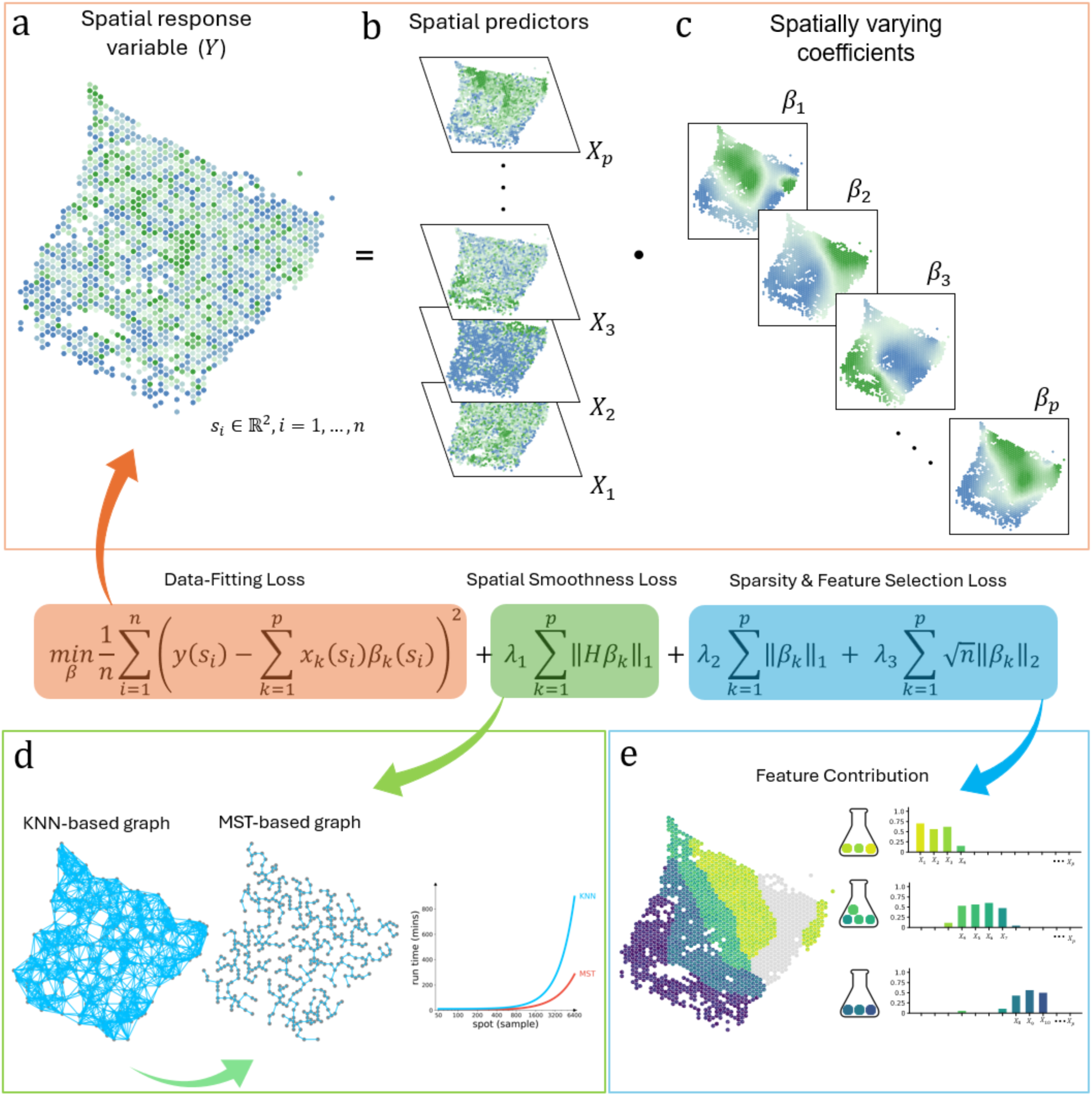
Workflow of the S3R method. (a-b) Input to S3R includes a spatial response feature 𝒚 and spatial predictor profile matrices 𝑿 across 𝒏 spatial locations. (c) The model outputs estimated spatially varying coefficients 𝜷_𝒌_(𝒔_𝒊_) for each predictor at each spatial location 𝒔_𝒊_**. Center, Composite loss function**: The estimation is performed by minimizing a composite loss function, which integrates three terms: a data-fitting loss, a spatial smoothness loss, and sparsity/feature-selection penalties. (d) A minimum spanning tree (MST) graph is constructed from spatial coordinates to enforce spatial coherence in coefficients 𝜷_𝒌_(𝒔_𝒊_). (e) The 𝑳_𝟏_penalty shrinks individual coefficients to zero, while the group 𝑳_𝟐_ penalty eliminates non-predictive features globally, collectively, they select predictors with spatially varying contributions to the response.

S3R is a flexible regression framework that accommodates diverse predictor–response configurations while estimating location-specific effects 𝜷_𝒌_(𝒔_𝒊_). For instance, when the response is the expression of a target gene and the predictors are the expressions of putative transcription factors, 𝜷_𝒌_(𝒔_𝒊_) quantifies the association of transcription factor 𝒌 with the target’s expression at location 𝒔_𝒊_. When the response is a gene’s expression and the predictors are spot-level cell-type proportions, 𝜷_𝒌_(𝒔_𝒊_) describes how local variation in the abundance of cell type 𝒌 relates to that gene’s expression across the tissue, thereby approximating the cell type–attributed expression of the target at location 𝒔_𝒊_. Finally, when the response is a target gene’s expression, and the predictors are assembled across levels, such as other genes (putative regulatory inputs), local neighborhood composition, and global niche context, the resulting coefficient fields 𝜷_𝒌_(𝒔_𝒊_) delineate how each level contributes to the target’s expression at location 𝒔_𝒊_ and how those contributions vary across space. In the sections that follow, we present case studies illustrating these settings and the breadth of biological questions S3R can address.

### Application and validation of S3R on recovering true target-TF regulatory relations on a DLPFC dataset

To demonstrate the ability of S3R on resolving spatially varying target–TF associations, we analyzed ST data collected from human dorsolateral prefrontal cortex (DLPFC) sections from Maynard et al. [26] (**Fig. 2a**). This dataset leverages well-defined laminar architecture and annotated layer-specific gene expression covering all six cortical layers and white matter (WM). We applied S3R to study the spatially varying regulatory relationships for a few selected target genes on two DLPFC slides (151673 and 151507; **Fig. 2a**). For each target gene, we modeled its expression as the response and used a curated panel of transcription factors as predictors; we refer to this fixed panel as “regulators” (**Fig. 2b-d**) and reserve “predictors” for the transcriptome-wide analysis below (**Fig. 2b–f**). The candidate transcription factors include all members from three Gene Ontology gene lists, including GO:0003700 (DNA-binding transcription factor activity), GO:0006355 (regulation of transcription, DNA-templated), and GO:0006351 (transcription, DNA-templated) [27]. Regulators or predictors expressed in less than 10% of spots of the corresponding sample are removed prior to analysis.

**Fig. 2.**
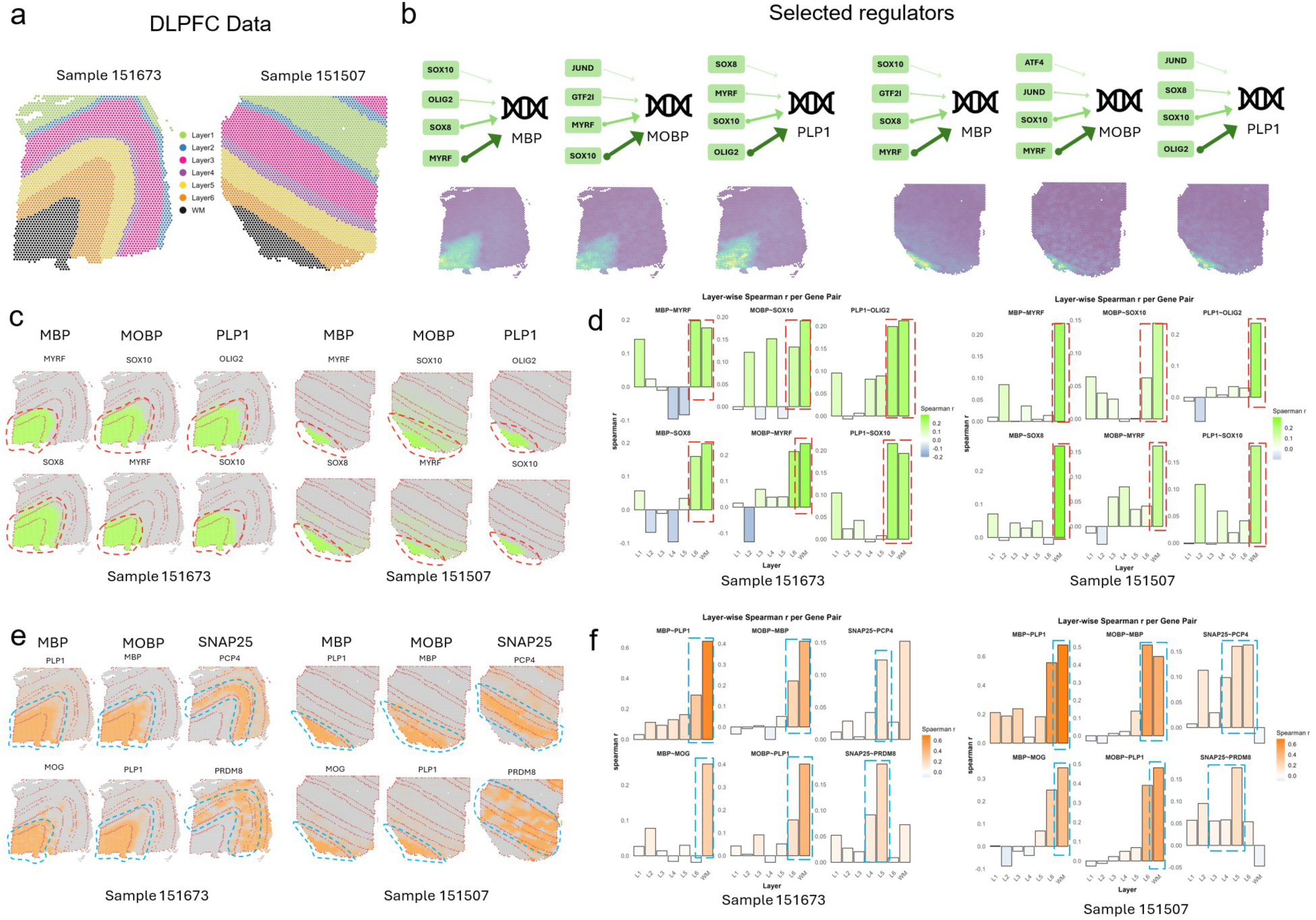
Computational evaluation of S3R on a brain DLPFC data. Two selected DLPFC sections analyzed in this study. (a) Top: S3R selected top regulators (green box) point to targets with arrow widths proportional to the inferred effect size (coefficient magnitude); Bottom: Spatial expression profiles of the target genes in the two DLPFC sections. (b) Spatial maps of the estimated coefficients for the top two S3R-selected regulators of three target genes, shown for both sections. Models here were fit using only a candidate set of ∼1,400 putative regulators. (c) Layer-wise correlations between the selected regulators and their targets using matched single-cell RNA-seq (*x-*axis: cortical layers; *y*-axis: correlation). Red dashed boxes mark layers with peak correlations. (d) As in c, but the model was fit using all genes. (e) As in d, but the model was fit using all genes.

We focused on three target genes with well-characterized white-matter-enriched expression: MBP, MOBP, and PLP1. After applying S3R, we selected regulators with non-zero location-specific coefficients, and obtained 26/16, 27/20 and 31/20 regulators for MBP, MOBP and PLP1 in the two sections respectively (selection procedure in Methods and Materials). The top four selected regulators for the three target genes in both slides are showcased in **Fig. 2b**, where the arrows connect each target gene to its top selected regulators by S3R among all the ∼1,400 candidates, with edge width proportional to the spatial effect size. **Fig 2c** shows the location-specific coefficients 𝜷_𝒌_(𝒔_𝒊_) for the three target genes and their top two selected regulators in the two DLPFC sections, and each figure panel corresponds to one target–regulator pair. The red dashed outlines highlight specific cortical layers where strong spatial effects per target were detected and revealed coherent laminar patterns: MBP showed associations with SOX8 and MYRF concentrated in WM across both sections; MOBP showed associations with SOX10 and MYRF prominent in WM with extension into layer 6; and PLP1 showed associations with SOX10 largely confined to WM, and with OLIG2 largely in WM extending to layer 6 (**Fig. 2c**).

For orthogonal validation, we analyzed matched single-cell RNA-seq data from the same cortical regions [28] (**Fig. 2d**). Here, we calculated the Spearman correlation of the target gene’s expression and all of the 1,400 candidate regulators’ expression in a layer specific manner. It turns out that top regulators selected by S3R show strong correlation with its target gene in the layer that is consistent with the S3R prediction (**Extended Figure 2a**). **Fig. 2d** provided a layer specific correlation between target gene and its selected top two regulators using the matched single cell RNA-seq data. The red dashed boxes indicate the layers where S3R selected regulators show the highest coefficients for the target, which coincide with the layers demonstrating the highest correlations among the regulator and target. This is consistent for all the three target genes and its top regulators across the two DLPFC samples.

To further demonstrate the S3R’s ability to deal with high dimensional feature space, we next extended the analysis to all expressed genes that have non-zero expression in at least 10% of the spots (**Fig. 2e–f**). This is in contrast to the analysis above where only transcription factors were included in the mode. In total, about 4,300 predictors were used for both sections. We here studied three genes as target genes, i.e., MBP, MOBP, and SNAP25. After applying S3R, we identified 29/16, 25/22 and 36/22 number of spatially varying and significantly associated predictors for MBP, MOBP and SNAP25, for the two slides. The estimated coefficients of the top two selected predictors, which associate with the respective response gene in a spatially varying manner, was showcased in **Fig. 2e**. The top predictors recovered by S3R again recapitulated known laminar organization: MBP was associated with PLP1 and MOG predominantly in WM; MOBP was associated with MBP and PLP1 in WM with partial extension into layer 6; and SNAP25 was associated with PCP4 and PRDM8 enriched in layer 5 and layers 3–5, respectively.

Similarly, we validated such layer-specific associations using matched single cell RNA-seq data. Again, the selected top layer specific predictors consistently show higher correlations with the target gene than the other non-selected candidate predictors (**Extended Figure 2b**). Also, layer-wise single-cell correlations peaked in the same layers highlighted by the spatial coefficient maps, corroborating the specificity of these associations estimated by S3R (**Fig. 2f**). Collectively, these results indicate that S3R accurately performs location-specific variable selection and recovers spatially coherent, biologically plausible target–TF associations from a curated TF panel and scales to transcriptome-wide predictors while preserving fine layer specificity across independent DLPFC sections.

### Application of S3R to recover cell type–specific expression and spatial gradients

The flexibility of S3R allows distinct spatial patterns to be interrogated by choosing the response and predictors in the regression model appropriately. To uncover cell type–specific spatial expression, we modeled, for each gene in turn, its spot-level expression as the response and the deconvolved cell-type proportions as predictors. This formulation is natural and useful for 10x Visium-or NanoString GeoMx DSP-based spatial transcriptomics data, where each spot/ROI contains multiple cells and the measured expression is a weighted average of constituent cell types. While most spatial deconvolution methods estimate only spot-level mixing proportions [29], in this formulation, the location-specific coefficients 𝜷_𝒌_(𝒔_𝒊_) quantify the association between cell type 𝐤 and the expression of the target gene at spatial location 𝐬_𝐢_, yielding a cell type–attributed coefficient field for each gene. We applied this framework to our in-house 10x Visium dataset from acute *H. ducreyi* infection of the skin [30] (**Fig. 3**). By assigning gene expression to specific cell types at each location, S3R could resolve where epithelial, stromal, and immune programs are deployed during the early host response to the bacteria infection, and further clarifies how local microenvironments orchestrate the acute response to *H. ducreyi* in situ.

**Fig. 3.**
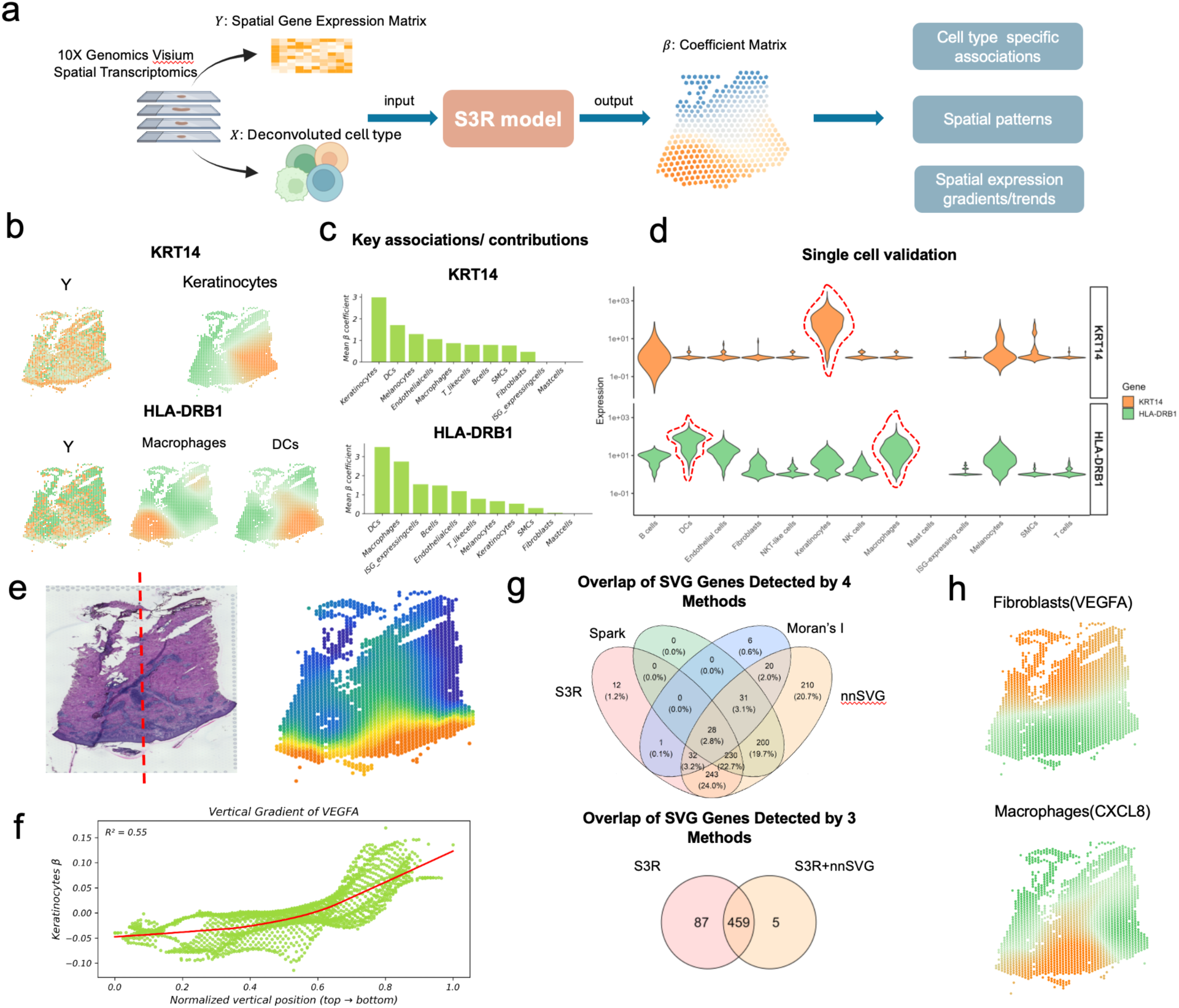
Application of S3R to spatial transcriptomics data from acute *H Ducreyi*-infected skin samples. (a) Inputs to S3R comprise the spot-level expression of a target gene (response) and spot-level cell-type proportions (such as deconvolved with RCTD) (predictors); the model outputs a spot-by–cell type coefficient matrix that approximates cell type–attributed, spatially varying expression for the target gene. (b) Spot-level expression (left) and S3R-estimated cell type–attributed expression maps (middle/right) for KRT14 (top) and HLA-DRB1 (bottom). (c) Mean cell type–attributed expression for KRT14 (top) and HLA-DRB1 (bottom) across spots (𝒙-axis: cell types; 𝒚-axis: mean attributed expression). (d) Distributions of KRT14 (top) and HLA-DRB1 (bottom) expression across cell types in matched single-cell RNA-seq. (e) H&E section of the slide (left) and the estimated spot specific expression of VEGFA by keratinocytes (right). The red dashed line indicates a normalized vertical coordinate perpendicular to the epidermis (bottom) and dermis (top) junction. (f) Relationship between normalized vertical position (𝒙-axis; epidermis to dermis) and keratinocytes-attributed VEGFA coefficients (𝒚-axis). (g) Overlap of spatially varying genes (SVGs) called by S3R, SPARK, Moran’s I, and nnSVG using spot-level expression (top) and using S3R-derived cell type–attributed expression (bottom). (h) Examples of gradients detected by S3R but missed by spot-level methods: fibroblasts–VEGFA (top) and macrophages–CXCL8 (bottom).

Genes expressed in at least 10% of spots were retained and analyzed. Predictors consist of RCTD-estimated spot-level proportions [29] for eleven cell types curated from matched single-cell data: keratinocytes, fibroblasts, melanocytes, smooth-muscle cells, endothelial cells, T/NK-like cells, macrophages, dendritic cells (myeloid and plasmacytoid combined to reduce collinearity), ISG-high cells, mast cells, and B cells (**Fig. 3a**). Exact thresholds and regularization parameters are provided in Methods.

As an initial validation, we examined two genes with well-established cell type specificity: KRT14, a structural keratin predominantly expressed by keratinocytes, and HLA-DRB1, a canonical antigen-presentation gene enriched in antigen-presenting cells. S3R recovered spatially coherent coefficient fields that attributed KRT14 to keratinocytes and HLA-DRB1 to macrophage and dendritic-cell compartments, with boundaries that mirrored histological features (**Fig. 3b**). A cell type was considered attributing to a gene’s expression when its coefficient field exhibited significantly non-zero values at certain spatial regions after spatial regularization (Methods). A downstream contribution analysis that ranks predictors by marginal effect confirmed keratinocytes as the dominant source attributed for KRT14 and macrophages/dendritic cells for HLA-DRB1 (**Fig. 3c**). These associations were corroborated in matched single-cell RNA-seq profiles, where expression localized to the expected cell populations (**Fig. 3d**).

A direct application of the resulting spatial and cell type specific expression from S3R is to examine the spatially varying expression gradients across different cell types. We next asked whether S3R resolves spatial gradient organization by using VEGFA as an example. We highlight VEGFA because it anchors angiogenic remodeling in acute cutaneous infection [31, 32], is robustly expressed at Visium resolution, has cell type–dependent sources in skin [33], and thus provides a test case of S3R’s ability to disentangle cell mixing and recover spatial gradients. In **Fig. 3e**, the red dashed-line in the H&E staining image marks the epidermal-dermal axis with epidermis and dermis towards the bottom/top of the image (left panel **Fig. 3e**), also called as normalized vertical coordinate. S3R attributed a prominent coefficient field to keratinocytes and fibroblasts. Specifically, keratinocytes-attributed VEGFA coefficients varied along the dermal–epidermal axis, and decreased toward deeper dermis and is the highest at the epidermis level (**Fig. 3e**). And the coefficient showed a strong monotonic trend with vertical coordinate (R² = 0.55, *P* < 1 × 10⁻¹⁵; **Fig. 3f**), whereas the raw spot-level VEGFA signal exhibited only a negligible association with the normalized vertical coordinate (R² = 0.0047, *P* = 1.7 × 10⁻⁴), illustrating how cell mixing can dilute gradients in aggregate signals.

Finally, we compared spatially varying genes (SVGs) identified by S3R with three widely used spot-level methods, namely Moran’s I [34], nnSVG [34, 35], and SPARK [34, 36], on the same section (**Fig. 3g**). S3R called SVGs based on the spatial and cell type specific expression out from S3R, combined with Moran’s I spatial autocorrelation (details in Methods and Materials). When applied directly to spot-level expression, cross-method concordance was modest: S3R, Moran’s I, nnSVG, and SPARK called 546, 118, 994, and 489 SVGs, respectively, with only partial overlap (**Fig. 3g**, top). The specifications of different methods were detailed in Methods and Materials. **Supplementary Table S1** listed the detailed SVGs called by each method. Overall, the consistency of the SVGs across different methods were poor; however, S3R detected SVGs overlap better with other methods (Top panel of **Fig. 3g**). For example, majority of SVGs called by nnSVG overlap with those by S3R; half SVGs detected by Moran’s I and Spark overlap with those by S3R. Because heterogeneous cell mixtures can drive method-specific calls, we reapplied nnSVG to the S3R-derived cell type–attributed expression matrices. Here, we didn’t apply SPARK as it only accepts count-based data as input. Removing mixture confounding in this way substantially increased concordance, yielding a larger shared set among the two approaches (**Fig. 3g**, bottom). As an example, S3R identified VEGFA attributed to fibroblasts and CXCL8 attributed to macrophages as SVGs. This means that fibroblast expressed VEGFA and macrophage expressed CXCL8 display spatial gradients, however, the spot-level expressions of VEGFA and CXCL8 didn’t display spatial gradients, and are missed by the spot-level SVG methods (**Fig. 3h**). This application illustrated an application of S3R on cell type–resolved modeling by revealing cell type specific expression, gradients and patterns that are diluted in aggregate signals.

### Application of S3R to recover cross–cell type, cross-gene four-way co-variation

Aside from recovering the cell type specific spatial expression gradients shown above, we here demonstrate another important utility where we used S3R to infer cell type–attributed, spatially varying gene fields and then quantified cross–cell type, cross-gene co-variation at spot resolution. This construction is motivated by the premise that intercellular crosstalk is expressed as coordinated, spatially localized deployment of gene programs across distinct cell types. By working with cell type–attributed coefficient fields rather than raw spot counts, we disentangle “who expresses” from “where it occurs,” allowing us to read out coupling between cell type pairs regarding gene programs. In doing so, the analysis moves beyond ligand–receptor lookups to a data-driven view of which cell types are coordinated, by which genes, and in which niches, thereby prioritizing concrete hypotheses for orthogonal validation (e.g., spatial proteomics, multiplexed IHC, perturbations) and clarifying how microenvironmental context organizes disease biology in situ.

For each gene *g*, we again modeled its spot-level expression as the response and deconvolved cell-type proportions as predictors, yielding location-specific coefficients 𝜷_𝒌,𝒈_(𝒔_𝒊_) that we interpret as the cell type–attributed expression profile for gene 𝐠 in cell type 𝐤 at spot 𝐬_𝐢_ (**Fig. 4a**). Stacking these coefficient fields across genes produces, for each cell type, a spots-by-genes matrix with many entries shrunk to zero by S3R’s sparsity penalties; additional post hoc filtering set small coefficients to zero when not significantly different from zero (see Methods). We applied this framework to our in-house 10x Visium sections from seven pancreatic ductal adenocarcinoma (PDAC) patients, and each sample contains cell types such as ductal cancer cells, fibroblast subtypes (permCAF and restCAF), immune populations, and endothelium [37].

**Fig. 4.**
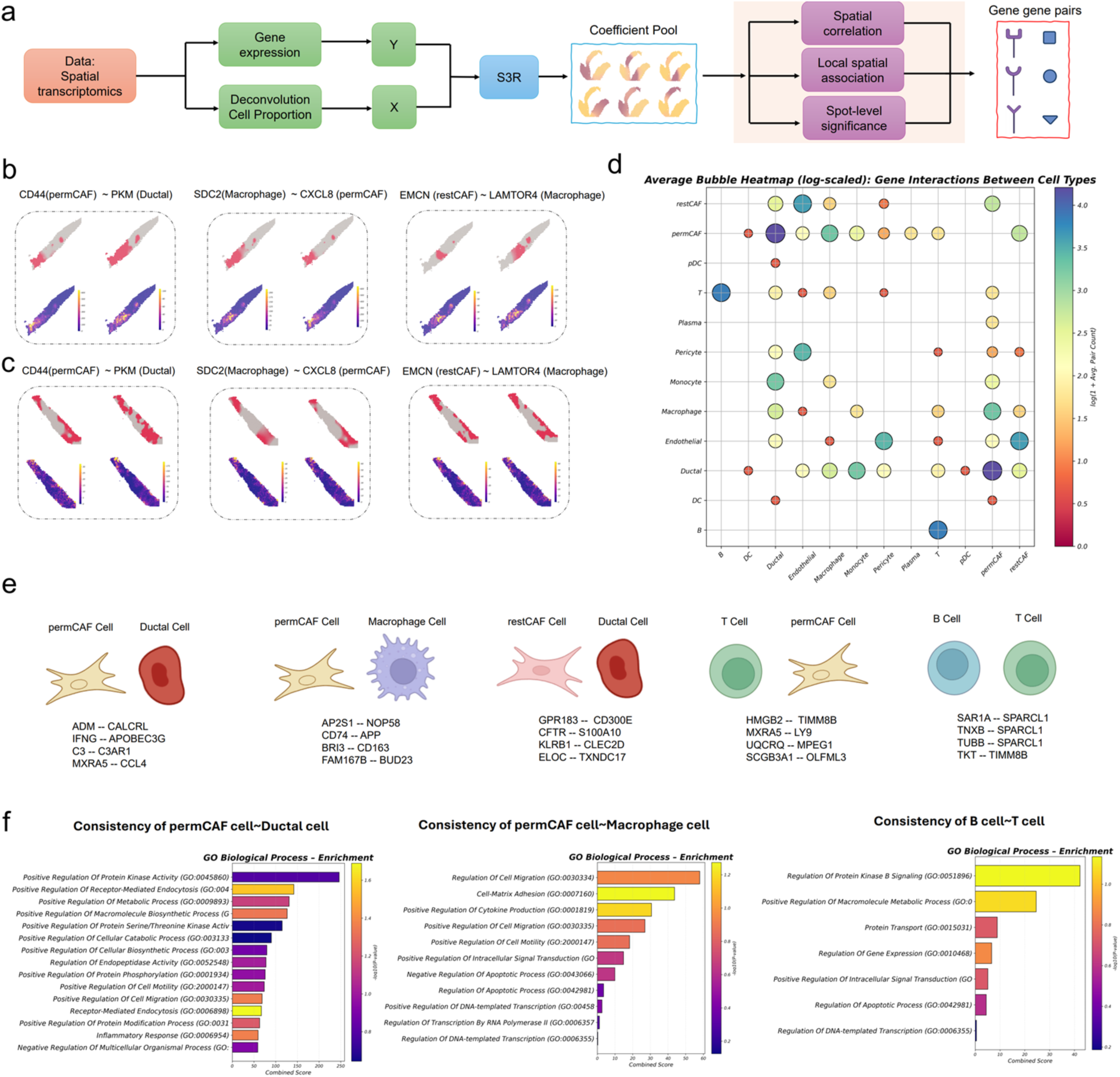
Application of S3R to PDAC spatial transcriptomics data. **(a)** Inputs to S3R comprise the spot-level expression of a target gene (response) and spot-level cell-type proportions deconvolved with RCTD (predictors); the model outputs a spot-by–cell type coefficient matrix for each gene. **(b)** Coefficient maps (top) for cell type pairs exhibiting co-variation between gene-gene pairs, and original spot-level expression of the same gene pairs (bottom), for three four-way associations in section 1. **(c)** Same as (b), but in section 2. **(d)** Bubble heatmap showing average interaction frequency/strength between cell-type pairs across seven PDAC sections. Bubble size and color represent interaction strength. **(e)** Representative top gene pairs for key cell type pairs (e.g., permCAFs–ductal, permCAFs–macrophages, B–T cells). **(f)** GO biological-process terms enriched by interacting genes for key cell type pairs.

To quantify cross–cell type gene co-variation, we considered any pair of cell types (𝐂_𝟏_, 𝑪_𝟐_) and any pair of genes (𝑮_𝟏_, 𝑮_𝟐_) and computed the Spearman correlation across spots between𝜷_𝑪𝟏,𝑮𝟏_(𝐬) and 𝜷_𝑪𝟐,𝑮𝟐_(𝐬). Because the 𝛃-fields are spatially smoothed, simple rank correlation approximated more elaborate spatial autocorrelation measures in practice; nonetheless, all significance assessments used sample-wise permutation controls and false-discovery-rate correction, and we retained only pairs that exceeded an effect-size threshold and replicated across sections (see Methods). The resulting quantities define a four-way tensor over genes and cell types that summarizes cross–cell type co-variation at tissue scale. **Supplementary Table S2** listed, for a pair of cell types, all the gene pairs that are significantly correlated after controlling for false discovery rate.

In **Fig. 4b-c**, we illustrated examples that highlight distinct spatial couplings that are consistent in two selected sections. A ductal–permCAF pairing showed coordinated spatial variation between ductal PKM and permCAF CD44, consistent with metabolic remodeling in the tumor epithelium aligned with CAF-mediated adhesion and matrix engagement (Left, **Fig. 4b-c**). A macrophage–permCAF pairing exhibited co-variation between macrophage SDC2 and permCAF CXCL8, in line with chemokine-rich stromal niches (Middle, **Fig. 4b-c**). And a restCAF–macrophage pairing displayed endomucin (EMCN) and LAMTOR4 co-variation suggestive of vascular–immune remodeling in CAF-adjacent regions (Right, **Fig. 4b-c**). In each case, the S3R coefficient maps (top row within each figure panel) localized a subset of the high-expression spots seen in the raw expression maps (bottom row within each figure panel), as expected when spot-level signals arise from mixtures and only a fraction of spots exhibit the gene in the cell type of interest.

We summarized cross–cell type coupling across the seven patient tumor slides by binarizing section-level associations into seven gene–gene–cell–cell tensors (1 = significant correlation, 0 = otherwise) and then aggregating across seven sections. The resulting average interaction strengths for a pair of cell types is quantified by the average counts of significant gene-gene pairs for the cell type pair. As displayed by a bubble plot in **Fig. 4d**, our analysis indicated prominent coupling between ductal cells and permCAFs, followed by restCAFs, monocytes/macrophages, and endothelium; macrophages coupled most strongly with ductal cells and permCAFs; and B cells showed their strongest coupling with T cells. We catalogued the top, consistently replicated gene pairs for key cell-type pairs (**Fig. 4e**). Pathway enrichment on genes participating in these cross–cell type couplings revealed kinase signaling, receptor-mediated endocytosis, and macromolecule biosynthesis in ductal–permCAF pairs (**Fig. 4f**, left), consistent with reports that bidirectional malignant cell–fibroblast crosstalk programs myCAF molecular/functional heterogeneity and promotes metastasis [38], that macropinocytosis sustains the myCAF state under glutamine stress to support ECM deposition and PDAC progression [39], and that TGF-β–driven myCAF differentiation rewires CAF metabolism toward glycolysis to supply tumor cells with metabolic intermediates [40–42]. Enrichment for cell migration/adhesion and cytokine production pathways in myCAF–macrophage pairs aligns with studies showing CAF programs that recruit and polarize myeloid cells and engage macrophage–CAF chemokine crosstalk [43, 44] (**Fig. 4f**, middle). Finally, enrichment for intracellular protein transport and signal transduction in B–T cell pairs is consistent with coordinated B–T activity within tumor-associated tertiary lymphoid structures linked to antitumor immunity and improved outcomes (**Fig. 4f**, right). **Supplementary Table S3** listed, for each cell type pair, the enrichment scores of Gene Ontology pathways and associated *p*-values for significantly correlated/interacting gene pairs.

### Application of S3R to high-resolution Xenium ST data

To demonstrate the utility of S3R on single-molecule, high-resolution ST data, we applied it to a 10x Genomics Xenium breast cancer dataset (**Fig. 5a**), which contains over 130,000 spatially resolved near-single cell resolution spots. Instead of modeling the entire slide, we focused on a subregion of approximately 40,000 cells (**Fig. 5b**), which represents a highly complicated tissue region with intermixed cell types, and **Fig. 5c** shows the relative proportions of annotated cell types within this selected region.

**Fig. 5.**
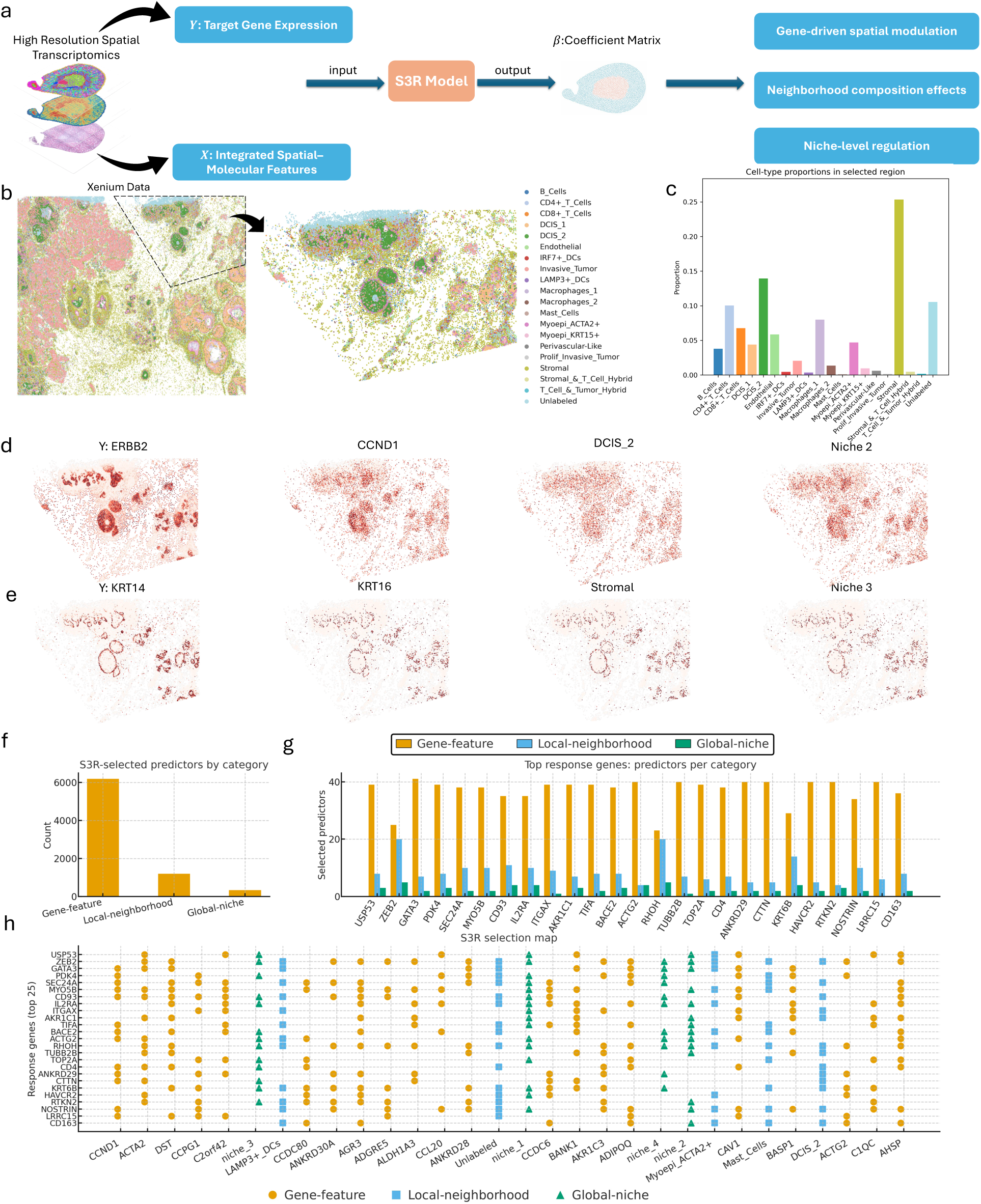
Application of S3R to 10x Xenium breast cancer spatial transcriptomics data. (a) Schematic overview of S3R applied to Xenium data. The model takes as input the spatial gene expression of each target gene (𝒀) and an integrated feature matrix (𝑿) combining other gene expressions, local neighborhood composition, and global niche-level representations. S3R outputs a spatially smooth and sparse coefficient matrix (𝜷) that captures gene-, local-, and global-level influences. (b) Xenium section showing cell-type composition map (left), and magnified view of the selected region (right). (c) The cell-type proportion distribution within the selected slide region. **(d-e)** Representative spatial maps of two example cases for ERBB2 and KRT14. (**d**) shows ERBB2 spatial expression (left) and the estimated spatial coefficients of its top selected predictors (CCND1, DCIS_2, Niche 2); (e) shows KRT14 spatial expression (left) and the estimated spatial coefficients of its selected predictors (KRT16, Stromal, Niche 3). (f) Category distribution of S3R-selected predictors in the Xenium breast cancer dataset, partitioned into gene-feature (intrinsic gene–gene effects), local-neighborhood (immediate cell-type/context features), and global-niche indicators. Bars show total selections per category. (g) The grouped bars quantify how many predictors were selected from each feature class (gene-feature, local-neighborhood, global-niche), for 25 selected genes. (h) S3R selection map for the 25 selected response genes (rows) versus top 30 predictors (columns). A mark indicates a selected predictor; marker shape encodes feature class.

We analyzed all 313 genes to delineate how spatial features across scales influence gene expression. Before fitting S3R, we constructed a joint feature matrix that explicitly represents the candidate sources of spatial influence for each gene, and assumed a gene’s spatial pattern can arise from three intertwined forces: (1) co-expression with other genes such as putative regulatory or pathway coupling, (2) the immediate cellular microenvironment, which is essentially the local neighborhood composition, and (3) the mesoscale niche in which a cell is embedded, which is the recurrent tissue contexts that extend beyond a local neighborhood. By placing these features side by side in the predictor matrix, S3R can disentangle their location-specific contributions and address questions that spot-or gene-centric analyses cannot: does a target’s expression track a local enrichment of particular cell types after controlling for other genes, do niche boundaries impose abrupt shifts in association strength, or are gene–gene relationships preserved uniformly or only within specific microenvironments.

Specifically, to construct the local neighborhood predictor in (2), for each cell, we identified its 30 nearest neighbors in Euclidean space, and computed the relative frequencies of different cell types to form a local composition vector, representing the immediate cellular microenvironment surrounding each cell. To capture a higher-level spatial “niche” structure in (3), we applied *k*-means clustering (*K* = 5) on all local composition vectors. The resulting clusters represent groups of cells that share similar local cellular composition profiles, and hence summarize recurring spatial microenvironments across the tissue. For example, in our analysis, Niche 1 represents epithelial–stromal transition zones with mixed epithelial, stromal, and immune components; Niche 2 corresponds to epithelial-or tumor-dominant dense regions; Niche 3 is enriched in stromal and myoepithelial cell types; Niche 4 consists of small endothelial or perivascular-like clusters; Niche 5 shows mixed immune–tumor regions, potentially reflecting immune–tumor interaction areas. The spatial local and global predictors, consistent for all the 313 target genes, are visualized in **Extended Fig. 3**.

Across each of the 313 target genes, S3R estimates a sparse and spatially smooth coefficient field 𝛽_*_(𝑠)that links 𝑌_*_to its predictors at location 𝑠. Small coefficients are shrunk to zero by the penalties and further removed by post-hoc filtering based on effect size and cluster-wise association tests (see Methods). The 𝜷-fields showed clear spatial organization that aligned with known tissue architecture. **Supplementary Table S4** listed the selected predictors for each of the target genes. Two illustrative examples of ERBB2 and KRT14 are shown in **Fig. 5c**: the 𝑌 panels display the raw target gene expression signal; the panels for selected features show the corresponding coefficient fields, highlighting gene-level, local neighbourhood, and global niche effects at cellular scale. The spatially resolved values of these S3R-selected features themselves is shown in **Extended Fig. 3**. In both cases, the selected variables align with established biology at gene, neighbourhood, and niche scales. For ERBB2 (**Fig. 5d**), S3R prioritized CCND1 (regulator of cell proliferation), a DCIS-enriched local neighbourhood (ductal epithelial context where HER2 activity concentrates), and Niche 2 (epithelial/tumour-dense), together consistent with HER2 (ERBB2) signaling coupled with CCND1 in compact epithelial territories [45]. For KRT14 (**Fig. 5e**), S3R highlighted KRT16 (frequently co-regulated with KRT14 by shared transcriptional programs), a stromal-rich local neighbourhood, and Niche 3 (stromal/myoepithelial-enriched). Together, this situates KRT14+ cells at ductal boundaries characterized by myoepithelial support and ECM remodeling.

We next summarized the overall findings across all 313 profiled genes by examining, for each target, the predictors selected by S3R. Of these, 214 genes had at least one predictor selected. Across 214 response genes, S3R identified 7,727 predictor selections spanning 314 unique predictors, partitioned into intrinsic gene–gene (gene-feature), local-neighborhood, and global-niche classes. The barplot provided overview (**Fig. 5f**) of the total selections of per class showing dominance of gene-level features with smaller yet consistent contributions from local cell-type context and broader niche indicators. We then picked 25 genes with the highest number of selected predictors, and showed the number of selected predictors across the three classes (**Fig. 5g**). And for the same 25 genes, we further mapped one-to-one predictor–target selections for the 30 most frequently selected predictors (**Fig. 5h**), to enabling direct comparison of class composition with the fine-grained selection map. Notably, the most frequently used predictors included a proliferative marker (CCND1), a myofibroblast/CAF program (ACTA2), immune context (LAMP3⁺ dendritic cells), and global niche domains (for example, niche_3), indicating that variation in target-gene expression is explained jointly by intrinsic programs, stromal remodeling, immune infiltration, and broader tissue context. Together, these patterns argue that S3R recovers interpretable predictors at three scales, within-cell programs, immediate cellular neighborhoods, and global niches; and that the balance of these scales is gene-specific, uncovering context dependencies that single-scale models would miss.

## Discussion

Here, we propose a spatially smooth and sparse regression model to fill a gap in spatial transcriptomics data analysis by estimating location-specific associations between a response feature and a high-dimensional set of candidate predictors while enforcing parsimony and spatial coherence. Conceptually, S3R treats “what relates to what” as a spatially varying problem: coefficients are learned as fields over the tissue, borrowing strength within neighborhoods yet permitting sharp transitions at histological boundaries. Methodologically, the combination of sparsity (𝑳_𝟏_ and group-𝑳_𝟐_) with a minimum-spanning-tree–guided smoothness term yields interpretable models that remain stable in the presence of noise, and collinearity, and computationally tractable at the scales typical of Visium and related platforms. A multi-GPU implementation and parallel hyperparameter tuning enable efficient training on large tissues and large predictor sets.

A central strength of S3R is its ability to operate in high-dimensional regimes without having to collapse to pre-screened, low-rank surrogates. In simulations, S3R recovered spatially varying coefficients, achieved accurate variable selection, and reproduced known spatial partitions. S3R’s regression formulation also makes it highly flexible to address distinct questions, because responses and predictors are user-defined. With a target gene expression as the response and other genes’ expressions (including TF), together with local and global contexts, as predictors, S3R estimates spatially varying contributions of gene intrinsic programs, as well as cellular neighbourhoods to a gene’s expression. With gene expression as the response and deconvolved cell-type proportions as predictors such as in Visium platform, S3R yields cell type–attributed expression fields for every gene, enabling gradient analyses and improved detection of spatially varying genes once cell mixing is mitigated. In addition, by correlating S3R coefficient fields across cell types and genes, one can construct cross–cell type, cross-gene co-variation tensors that prioritize candidate interactions for follow-up. This analysis complements existing computational approaches to cell–cell communication, where single-cell RNA-seq loses spatial coordinates and therefore cannot directly assess proximity-dependent effects [46–48], and spot-level spatial methods may lack cell-type specificity when mixtures are present [3–6]. Operating on S3R’s cell type–attributed coefficient fields resolves both issues, by preserving spatial localization and attributing signal to specific cell types and genes. Importantly, it is not restricted to ligand–receptor pairs but to alternative mechanisms encompassing all possible genes.

In summary, S3R provides a general, scalable approach for modeling spatially varying relationships in high-dimensional spatial transcriptomics data. We anticipate S3R will serve as a practical tool for a broad class of spatial analyses and as a bridge from descriptive maps to mechanistic investigation of tissue microenvironments. Despite these promising results, some limitations remain. S3R estimates associations, not causal effects; interpreting coefficient fields mechanistically requires orthogonal evidence such as spatial proteomics and perturbations experiments. The accuracy of cell type–attributed fields depends on the quality of deconvolution, and uncertainty propagation from deconvolution to S3R would make downstream inferences more rigorous. Like any smoothing approach, there is a bias–sharpness trade-off. While the MST-guided term preserves abrupt changes, automatic boundary detection and adaptive, data-driven neighborhoods could further improve edge preservation. Finally, formal inference on the coefficient fields and integration with additional modalities are represent future extensions.

## Methods

### The motivation of S3R model

Because conventional regression ignores spatial structure and classical spatial models cannot accommodate the high-dimensional predictors typical of ST data, spatial transcriptomics sits at a methodological gap between spatial statistics and high-dimensional regression [1, 49, 50]. One line of work addresses spatial homogeneity by allowing regression coefficients to vary over space, treating them as smooth functions. Spatially varying-coefficient models often use Gaussian-process fields or conditional autoregressive priors to encourage spatial coherence, and related non-Bayesian approaches include geographically weighted regression [49, 51–53]. These models capture gradual spatial trends in covariate effects, for example, a gene’s influence from a tumor core to its periphery, but purely smooth formulations have important limitations for ST analysis. In practice, tissues contain discrete microenvironments where effects shift abruptly; modeling such effects typically requires sparsity-and edge-preserving which smooth-only models do not readily provide.

Another class of methods addresses spatial heterogeneity by imposing penalized regression frameworks that combine sparsity and spatial structure. A prominent example is the fused lasso, which adds an 𝑳_𝟏_penalty not only on the regression coefficients (as in the ordinary lasso) but also on differences between coefficients in neighboring spatial locations [54]. This encourages solutions that are sparse and also locally coherent in space [54]. To perform variable selection in high dimensions, the group lasso is a useful regularization approach that selects or drops predefined groups of features as a whole [55]. In a spatial context, one can define each gene’s coefficient profile across all locations as a group. A group-lasso penalty then either retains that gene by allowing its coefficients to be nonzero at some locations, or sets the entire coefficient surface to zero, thereby performing gene selection. Recent methods in high-dimensional spatial regression have begun to integrate these ideas. Notably, Li and Sang (2019) proposed a spatial homogeneity pursuit method (SCC) that encourages spatial similarity in regression coefficients between locations connected by an Minimum Spanning Tree, to automatically detect cluster patterns and change points in coefficients [56]. Building on this concept, Zhong et al. (2023) introduced the Regularized Spatially Clustered Coefficient (RSCC) model to address large spatial datasets with many predictors [18]. RSCC combines a MST-guided graph penalty with a group lasso penalty to simultaneously achieve variable selection and spatially clustered coefficient patterns [18]. These developments underscore the value of penalties that accommodate abrupt jumps in coefficients while selecting relevant variables. However, their primary focus is on partitioning the space into homogeneous clusters; fully leveraging continuous gradients in the estimated coefficients, such as in the case of disease microenvironment of spatial transcriptomics data, remains underexplored. Moreover, while the group-lasso component in RSCC promotes group-level interpretability but does not yield location-specific sparsity, that is, once a predictor (or group) is selected by RSCC, it tends to be retained across all locations. An individualized and hence location-specific sparsity would further enhance biological interpretability, but the resulting computational burden demands scalable optimization. Overall, there remains a need for methods that adaptively balance smooth and discontinuous spatial variation in covariate effects, enable spot-level variable selection, and remain computationally feasible in high dimensions. Here we present Spatially Smooth and Sparse Regression (S3R) to address these challenges with a new regularization scheme that *jointly* penalizes redundant coefficient values and large differences between neighboring spatial coefficients.

### The overall model framework of S3R

Suppose a set of high dimensional spatial variables (𝑋(𝑠_i_), 𝑦(𝑠_i_), 𝑖 = 1,…, 𝑛), are observed at locations 𝑠_1_,…, 𝑠_n_ ∈ ℝ^2^, where the response variable 𝑦(𝑠_i_) is assumed to be spatially correlated, and 𝑋(𝑠_i_) = (𝑥_1_(𝑠_i_),…,𝑥_p_(𝑠_i_))*^T^* is the *p*-dimensional vector of explanatory variables for the observation located at 𝑠i. Consider a spatial regression model with spatially varying regression coefficients

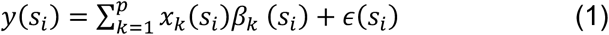

In our spatial transcriptomics dataset, we could observe only one sample at each location, making the model (1) ill-posed if without any restrictions on 𝛽_1_(𝑠_1_). For example, with only one spatial realization (𝑋(𝑠_1_), 𝑦(𝑠_1_), 𝑖 = 1,…, 𝑛) at *n* observed locations, the regression model (1) can be written in the matrix form

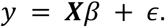

Here, we stack all the coefficients into a vector 𝛽 ∈ 𝑅^pn^, 𝛽 =(𝛽_1_(𝑠_1_),…,𝛽_1_(𝑠_n_),…, 𝛽_p_(𝑠_1_),…,𝛽_p_(𝑠_n_))^T^ and the design matrix 𝑿 = [𝑑𝑖𝑎𝑔(𝒙_1_),…,𝑑𝑖𝑎𝑔(𝒙_p_)] is an 𝑛 × 𝑛𝑝 matrix with 𝑿_k_ = (𝑥_k_ (𝑠_1_), 𝑥_k_ (𝑠_n_))^T^. Clearly, this regression problem needs to be regularized since there are more variables than observations. For spatial problems, it is expected that association between a response variable and explanatory variables at nearby locations is highly likely to be homogeneous. In addition, the number of candidate explanatory variables can be large, and we need to select the ones that are truly associated with the response variable. This motivates us to assign a few regularization functions for 𝜷 reflecting such spatial coherence and feature sparsity patterns.

Specifically, we propose to estimate 𝜷 by minimizing the following objective function

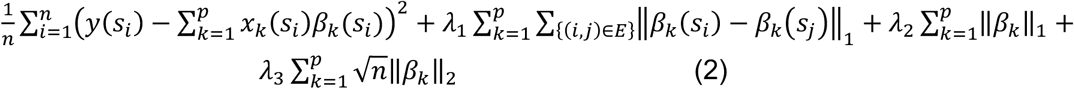

Here, 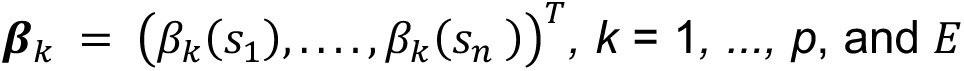 is the edge set of a graph consisting of *n* vertices, where each vertex corresponds to one observed spatial location, and each edge connects two connected locations. 𝜆_1_,𝜆_2_, 𝜆_3_ are tuning parameters determining the strength of penalization of spatial smoothness, individual sparsity and group sparsity. Specifically, the term associated with 𝜆_1_ is to encourage homogeneity between two regression coefficients if their corresponding locations 𝑠_i_ and 𝑠_j_ are connected by an edge in 𝑬. The term associated with 𝜆_2_ induces element-wise sparsity such that each location may have a different non-zero coefficients; 𝜆_3_ promotes group sparsity at the predictor level such that one predictor may be removed globally. For simplicity, we reformulate (2) as follows:

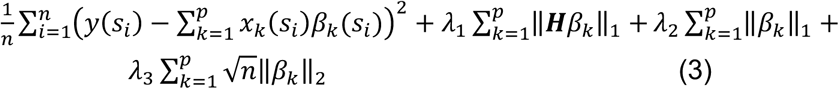

where 𝑯 is an 𝑚 × 𝑛 matrix constructed from the edge set 𝐸 with 𝑚 edges, similar to [56]. For an edge connecting two locations 𝑠_i_ and 𝑠_j_, we represent the penalty term 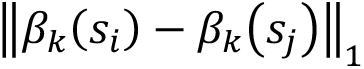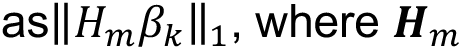 is a row vector of 𝑯 and contains only two nonzero elements, 1 at the 𝑖 th index and −1 at the 𝑗 th.

### The spatial smoothness control

To construct the edge set 𝑬, for most methods, the graphs are constructed by *k*-nearest neighbors or neighbors within a certain radius such that all points are connected, in which case, the number of edges is greater than the number of nodes. While many algorithms have been developed to solve the generalized lasso problem, for an arbitrary 𝑯 with *m > n* for large *m* and *n*, these algorithms are computationally very costly. An appropriate choice for 𝐸 that balances model accuracy and computational efficiency should only include coordinate pairs close to each other, should lead to connectivity of all data points, and should have no redundant pairs. One choice of 𝐸 that satisfies all three of these criteria is the edge set of a Minimum Spanning Tree (MST), similar to SCC and RSCC [18, 56].

Given an undirected graph 𝐺 = (𝑉, 𝐸_<_) with a weight function 𝑑(𝑒) that assigns a weight to each edge 𝑒 in an edge set 𝐸_<_, an MST is defined as the subgraph 𝑇 = (𝑉, 𝐸), 𝐸 ⊆ 𝐸_<_that connects all vertices without any cycles and minimizes ∑_e∈E_ 𝑑(𝑒).It is known that an MST has |𝑉| vertices and |𝑉 − 1| edges. It is known that for any cut of a given connected graph 𝐺, the minimum-weight edge that crosses the cut is in the MST for 𝐺. In our implementation, taking a given set of *n* spatial locations as vertices and the Euclidean distances between locations as the edge weights, we by default construct a *k*-nearest neighbors (KNN) graph (e.g., 𝑘 = 7) for small-to-moderate datasets (𝑛 ≤ 1000), while allowing users to choose either KNN or an MST. For larger datasets (𝑛 > 1000), we by default construct a Minimum Spanning Tree (MST), 𝑇 = (𝑉, 𝐸), consisting of 𝑛 − 1 edges with all *n* locations connected. By the construction, 𝑇 provides a connected, acyclic graph that compactly represents the spatial topology of the 𝑛 observed points and is thus an appropriate choice satisfying the aforementioned desired properties for edge set selections. In this case, 𝑯 is a full row rank matrix encoding the 𝑛 − 1 penalty terms on the coefficients 𝛽_k_.

### The algorithm implementation

We optimized the algorithm using Adam [23], combined with cosine/step-wise learning rate adjustments, gradient clipping, and early stopping to ensure stable convergence during training. The entire implementation uses batch mode and fully utilizes GPU acceleration, allowing the model to run on large datasets. Specifically, to allow efficient use of GPU: 1). When computing the first loss term, we performed element-by-element multiplications and then sum the results by feature, avoiding the time-consuming computation of large matrices; 2). When computing the graph-fused lasso loss, we prepare the edge indices and perform the computation on the GPU using an aggregation/difference operation, which results in a computational complexity of 𝑶(𝒎𝒑), linear in the number of edges 𝒎 and the number of features 𝒑; 3). When computing the group sparsity term, we rely on PyTorch autograd’s subgradient implementation and add a small 𝝐 stabilization term to prevent instability near zero.

### Model and hyperparameter selection

Our model has three hyperparameters: 𝜆, that controls the spatial smoothness of the regression coefficients; 𝜆. that: controls sparsity of individual parameters; and 𝜆: that controls group-level sparsity. These hyperparameters are selected by cross validation. Instead of randomly splitting the spots into training and testing sets, we divided the spatial region into several spatially coherent blocks, using some blocks for training and the rest for validation. This ensures that the model can generalize in space. Because we need to tune 𝜆_1_,𝜆_2_,𝜆_3_ together, the number of possible combinations is very large. If we use brute-force grid search or pure random search, the computation would be too expensive. To solve this, we use Optuna [25]., an automatic hyperparameter optimization tool. It searches hyperparameter combinations in a random + intelligent way within given ranges. Specifically, 𝜆_1_,𝜆_2_,𝜆_3_ are sampled on a log scale, namely, 𝑙𝑜𝑔𝑢𝑛𝑖𝑓𝑜𝑟𝑚(1𝑒 − 6, 1𝑒2)). In Optuna, each attempt of one hyperparameter combination is called a trial. We start many trials at the same time, each trial corresponds to a different hyperparameter set. These trials are assigned to different GPUs to run in parallel, so we can test many combinations at the same time and make the search much faster.

### Post-hoc significance evaluation of the coefficients

Inference of high dimensional regression is challenging, and even more in our case with three different levels of penalty functions. We implemented a post-hoc variable selection procedure to further select informative predictors with significantly non-zero coefficients. We borrow the idea in linear regression with lasso, where Lasso will be run first, and then the retained variables, much less than the original set, will be fit into the linear regression model, and inference of coefficients could be conducted and those non-significantly different from zero will be removed. Here, to achieve spatially varying variable selection, we conduct a model refitting for each spot using linear regression by using the spot and its neighbouring spots, by assuming that the neighboring spots share the same coefficient scales. To speed up the computation, in practice, we dissect the spatial map into multiple clusters for the linear regression model fitting. A coefficient was considered significant for the spot, if the associated 𝑝-value for the predictor in the linear regression is below the threshold (adjusted 𝑝 < 0.05).

### Deducing cell type specific and spot-level expressions

We applied RCTD for spatial deconvolution [29]. In terms of single cell expression reference and annotation required by RCTD, for the skin infection dataset, we utilized matching single cell RNA-seq data from the same study [30]; for the PDAC dataset, we utilized matching single cell RNA-seq data from the same group [37]. By regressing a gene’s expression on the cell type proportions using S3R, we deduced cell type–specific and spot-level expression. The resulting coefficient field 𝛽_g,c_(𝑠_i_) quantifies how strongly a given cell type explains the expression of the target gene at each spatial location. Here, we applied a second post-hoc parameter shrinkage additional to the procedure mentioned above: 1) only non-negative coefficients are retained and all negative-coefficients are shrunken to zero; 2) positive coefficients are only allowed in spots where the cell type has non-zero abundance. This two-step selection procedure is implemented because the cell types could only contribute to a gene’s expression positively; and if the cell type is not present, it should not contribute to the expression of the genes. In this way, the coefficients provide two layers of information: (i) cell type specificity, because predictors correspond to deconvolved cell-type proportions, and (ii) spot-level resolution, because coefficients vary across tissue locations. The resulting cell type by spots coefficient matrices could be utilized for downstream analysis such as spatially varying genes detection, and cross-cell and cross-gene interaction.

### Detecting spatially varying genes (SVG)

**S3R**: **S3R** adopts a multi-step procedure in detecting spatially variable genes. Based on S3R output of the coefficients by regressing a gene’s expression on the cell type proportions, we first conduct a post-hoc variable selection for all the spots, which will further shrink many coefficients to be zero. For each gene, we looked at its spot by cell type coefficient matrix. Then we used Moran’s I to test spatial variation. In other words, for each gene–cell type pair, we checked if its coefficients showed spatial structure.

**Moran’s I**: For each gene, Moran’s I is computed on expression profile across spots, and significance is assessed by permutation. A significantly positive Moran’s I indicates the presence of non-random spatial structure.

**nnSVG: nnSVG** (**n**earest-**n**eighbor **S**patially **V**arying **G**enes) models gene expression using a nearest-neighbor Gaussian process approximation. Expression is decomposed into non-spatial and spatially structured components. A gene is designated as an SVG if the estimated spatial variance is significantly greater than zero. In our analysis, nnSVG was implemented using the nnSVG R package, with expression counts and spatial coordinates encapsulated in a SpatialExperiment object. We used default settings with multi-threading enabled (n_threads = detectCores() – 1), and retained genes with adjusted 𝑝 < 0.05 as SVGs.

**Spark: Spark** (**S**patial **PA**ttern **R**ecognition via **K**ernels) employs generalized linear spatial models with multiple kernel functions to capture diverse forms of spatial dependence, including local hotspots, broad gradients, and periodic structures. For each gene, kernel-specific spatial effects are jointly tested, and 𝑝-values are combined to robustly detect spatially variable expression across multiple spatial scales. We applied SPARK using the official R implementation, providing raw count matrices and spot coordinates as input. Variance components were estimated with spark.vc and significance was assessed with spark.test under default settings. Genes with adjusted 𝑝 < 0.05 were called as SVGs.

### Pathway enrichment analysis

We performed Gene Ontology enrichment with the R package enrichR [57]. Due to our stringent control of the detected interacting gene pairs for each cell type pair, the number of interacting genes is of small scale. Hence, standard multiple-testing correction of enrichment analysis is underpowered and can mask coherent biology. Accordingly, we report raw p values together with enrichment scores, and interpret them as exploratory indicators of pathway coherence rather than confirmatory hypothesis tests.

### Datasets used

Both publicly available and in-house generated spatial transcriptomics datasets were used for benchmarking and case studies in this work. DLPFC datasets was obtained publicly from Maynard et al [26] (visium ST data) and [28] (matching single cell RNA-seq data); the *H. Ducreyi* infected skin dataset, both ST and scRNA-seq data, was an in-house dataset previously reported at Brothwell et al [30]; the UNC PDAC dataset was another in-house dataset previously reported at [37]; the Xenium based breast cancer dataset was obtained from 10xgenomics.com.

### Computational resources

All computations were performed on the Big Red 200 supercomputer at Indiana University, an HPE Cray EX system equipped with GPU-accelerated nodes (each with four NVIDIA A100 GPUs, AMD EPYC 7713 processors, and 512 GB memory). For a dataset of size 𝑁 = 3050 (spatial locations) and 𝑝 = 10,986 (genes), the complete pipeline required about 4 hours on a single GPU node (using 4×A100 GPUs, 16 CPU cores, and 32 GB memory per task). This runtime covers both the hyperparameter optimization using Optuna and the model training. Once the optimal hyperparameters were determined, the final estimation of the coefficient matrix could be completed within only a few minutes.

## Code availability

All codes to implement the S3R model, as well as to reproduce the results across the work, is available via GitHub at https://github.com/ZhouXY199502/S3R.

## Acknowledgments

This work was supported by research grants IIS-2145314 and DBI-2047631 from the National Science Foundation; and R35GM155028 from the National Institutes of Health; and RSG-24-1321371-01-CDP from the American Cancer Society; and the OHSU Brenden-Colson Center for Pancreatic Care.

## Author contributions

C.S. conceived the methodological framework. Z.C. designed all the real-world data applications. Z.X. designed the algorithm, simulation validation, and conducted the case studies.

D.P. provided key guidance on algorithm implementation. X.W., N.B. and Z.N. contributed to methodological guidance and discussion interpretation. P.L.X. and J.J.Y. provided well-annotated PDAC spatial and single cell RNA-seq data. P.L.X., J.J.Y., S.R and Z.T. contributed to applications of methods to PDAC. Z.C., Z.X., C.S. wrote the manuscript.

## Competing interests

The authors declare no competing interests.

## Supplementary

### Evaluation of S3R on simulation data

To rigorously validate the efficacy of the S3R method, we conducted a systematic evaluation using synthetic datasets. Ground-truth coefficient matrices were generated with 3–5 distinct spatial clusters and varying feature sparsity levels to reflect biologically plausible regulatory networks. Additive Gaussian noise (*σ*=0.5,1,2) was introduced to simulate technical variability, and feature dimensionalities spanned 20, 100, and 1000, representing low to high-dimensional feature spaces. A visual comparison of the clustering accuracy between S3R and four other competing methods, including RSCC, EN, GWL and VS-GPSVC was shown in **Extended Fig. 1a**. In this simulation scenario for the left four columns, the predictor matrix 𝑿 is of dimension 100×20, depicting a relatively low-dimensional feature setting; the predictor matrix 𝑿 is of dimension 100×2000 for the right two columns, depicting a high dimensional setting. We introduced different number of coefficient clusters that exist among the response and the predictors, such as 3,4,5,6. The detected clusters by different methods were colored differently. The spatial map on the first row shows the true coefficient matrix. The map in the second row represents the coefficient clusters given by our proposed S3R algorithm, which shows a high degree of similarity with the true spatial dissection, similar for the other methods shown in the rest of the rows.

A more detailed quantitative evaluation is presented in Extended **Fig. 1b**, where we report the Adjusted Rand Index (ARI) between the true coefficient clusters, and the coefficient clusters given by different methods, obtained under five cluster scenarios (3–6 clusters) and three noise levels (σ = 0.5, 1, 2). The first four columns represent the low-dimensional settings (N × p = 100 × 20 to 2000 × 20), while the last two illustrate the high-dimensional cases (N × p = 200 × 1000 and 400 × 1000). Across all panels, ARI scores decline as the Gaussian noise increases, yet S3R consistently attains the highest values, outperforming RSCC, EN, GWL, and VS-GPSVC in every configuration. Notably, the gap between S3R and the competing methods widens in the high-dimensional regime, underscoring the effectiveness of its sparsity and spatial smoothness constraints when 𝑝 far exceeds 𝑁.

### Runtime comparison of S3R and competing methods

To assess computational scalability, we compared the runtime of S3R with RSCC, Elastic Net (EN), Graph-guided Weighted Lasso (GWL), and VS-GPSVC across datasets with varying sample sizes (N = 200–2000) and feature dimensions (P = 20–1000). As shown in Supplementary **Fig. 1**, all methods perform similarly in low-dimensional settings (e.g., N = 200, P = 20), although GWL consistently incurs higher runtime. With increasing dimensionality (e.g., N = 400, P = 100; N = 1000, P = 100), VS-GPSVC becomes substantially slower. In high-dimensional regimes (e.g., N = 200–2000, P = 1000), S3R maintains practical runtime while several competing methods either slow down considerably or fail to complete (red “X” marks). Across all scenarios, S3R achieves favorable runtime relative to accuracy. All runtime experiments were conducted on a personal PC.

---

**Supplementary Figure 1.**
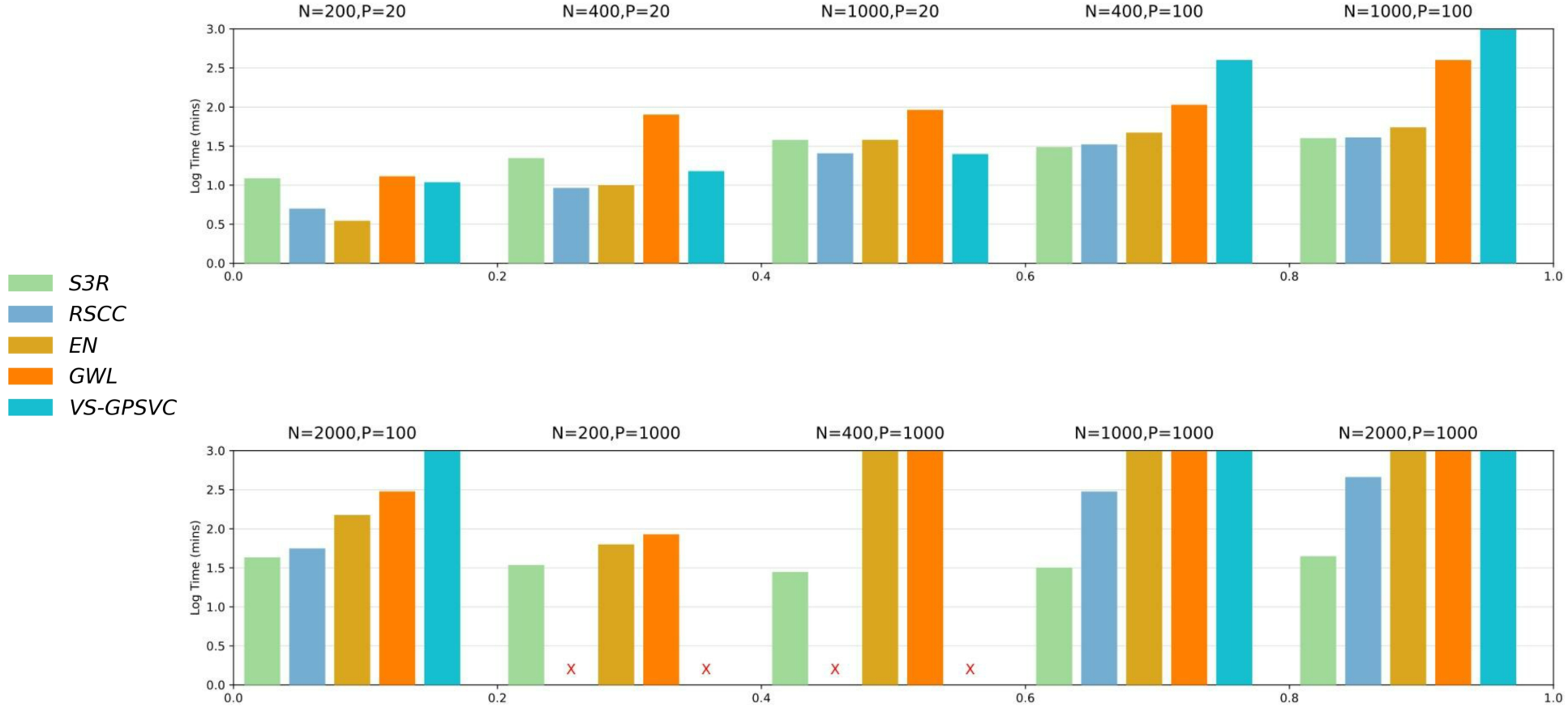
The runtime comparison of S3R and other methods. Supplementary Tables

Supplementary Table S1: SVG genes detected by 4 methods.

Supplementary Table S2: For each cell type pair, the list of S3R detected interacting gene-gene pairs across all seven PDAC datasets.

Supplementary Table S3: For each cell type pair, the enriched Gene Ontology pathways and enrichment p-values using gene pairs that are detected by S3R to be interacting for the cell type pair.

Supplementary Table S4: The selected gene-, local-and global-level features for each of the 313 target genes.

**Extended Figure 1.**
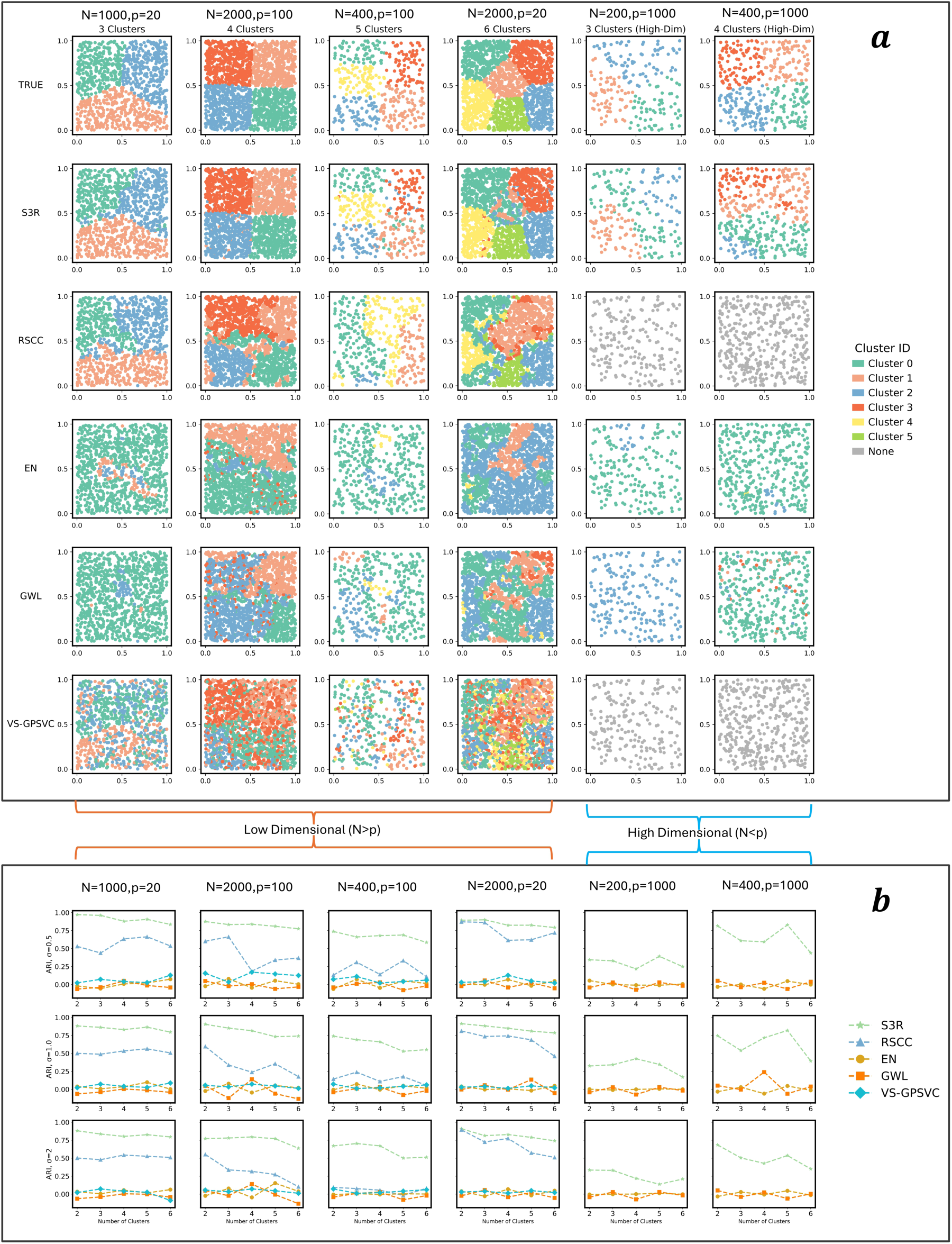
Evaluation of S3R on simulation data. a, A visual comparison of S3R with other benchmarking methods in recovering the coefficient clustering patterns across various simulation scenarios. From left to right are different sample size (𝑵) and feature dimensions (𝒑). From top to bottom are ground-truth coefficient cluster patterns, and estimated coefficient clusters by different methods. 𝒙-axis and 𝒚-axis are spatial coordinates. Colors of the points indicate clusters of coefficients given by different methods. **b,** Quantitative evaluation of S3R with other competing methods based on Adjusted Rand Index (ARI) of the estimated coefficient clusters with ground truth, under various simulation settings. The dashed line indicates different methods. 𝒙-axis shows different numbers of simulated coefficient cluster patterns, and 𝒚-axis shows the ARI. From left to right: different sample size (𝑵) and feature dimensions (𝒑).

**Extended Fig. 2.**
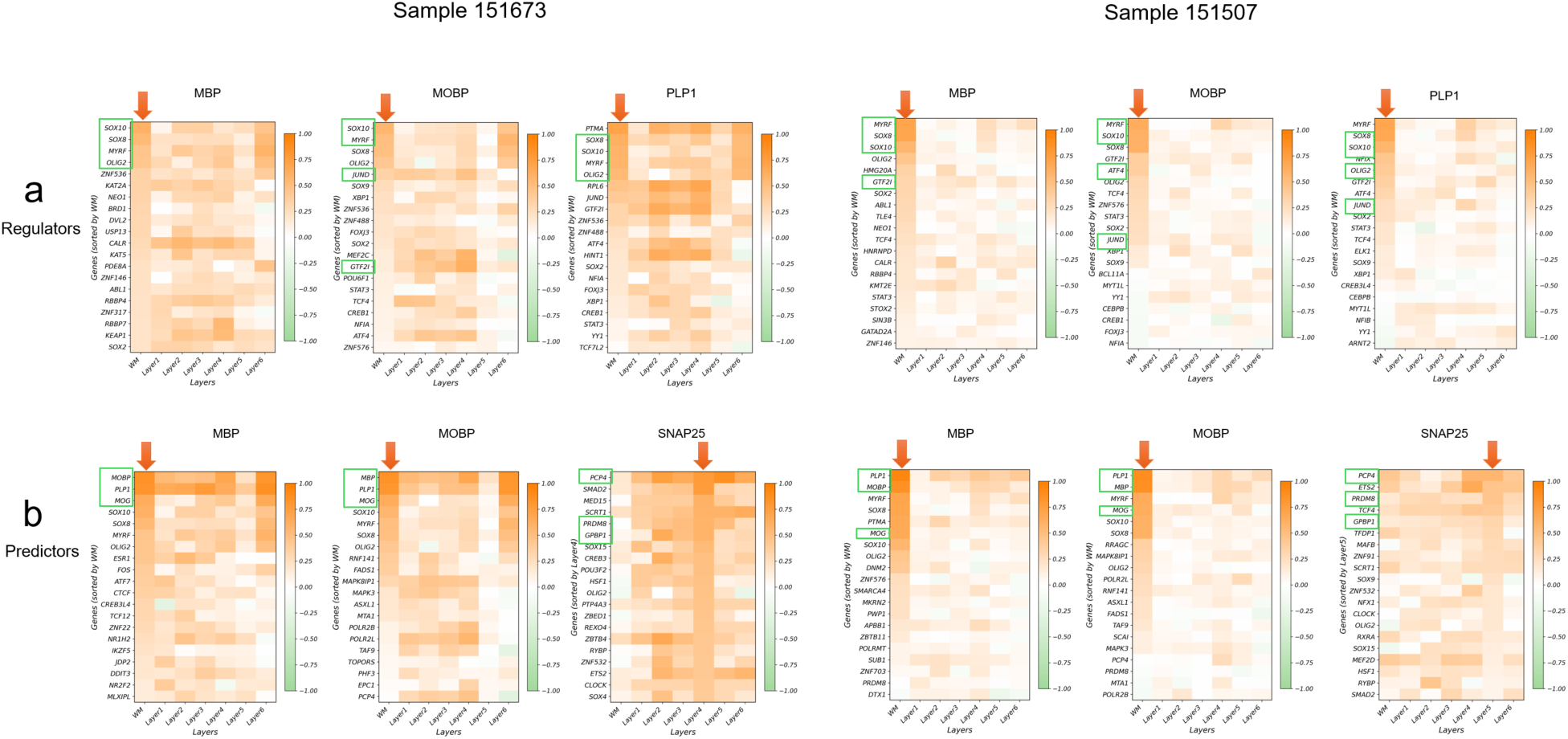
Layer-wise Spearman correlations between each target gene and S3R-selected regulators/predictors for the two DLPFC sections, using matched single cell data of DLPFC. a) shows correlations with regulators and b) shows correlations with predictors that encompass all genes. Genes highlighted with green boxes are the top candidates used in Fig. 2 (four regulators and three predictors). 𝑥-axis shows different layers, and 𝑦-axis shows different genes ranked by the average coefficient magnitude estimated by S3R. This figure provides a quantitative, layer-specific complement to Fig. 2d,f.

**Extended Fig. 3.**
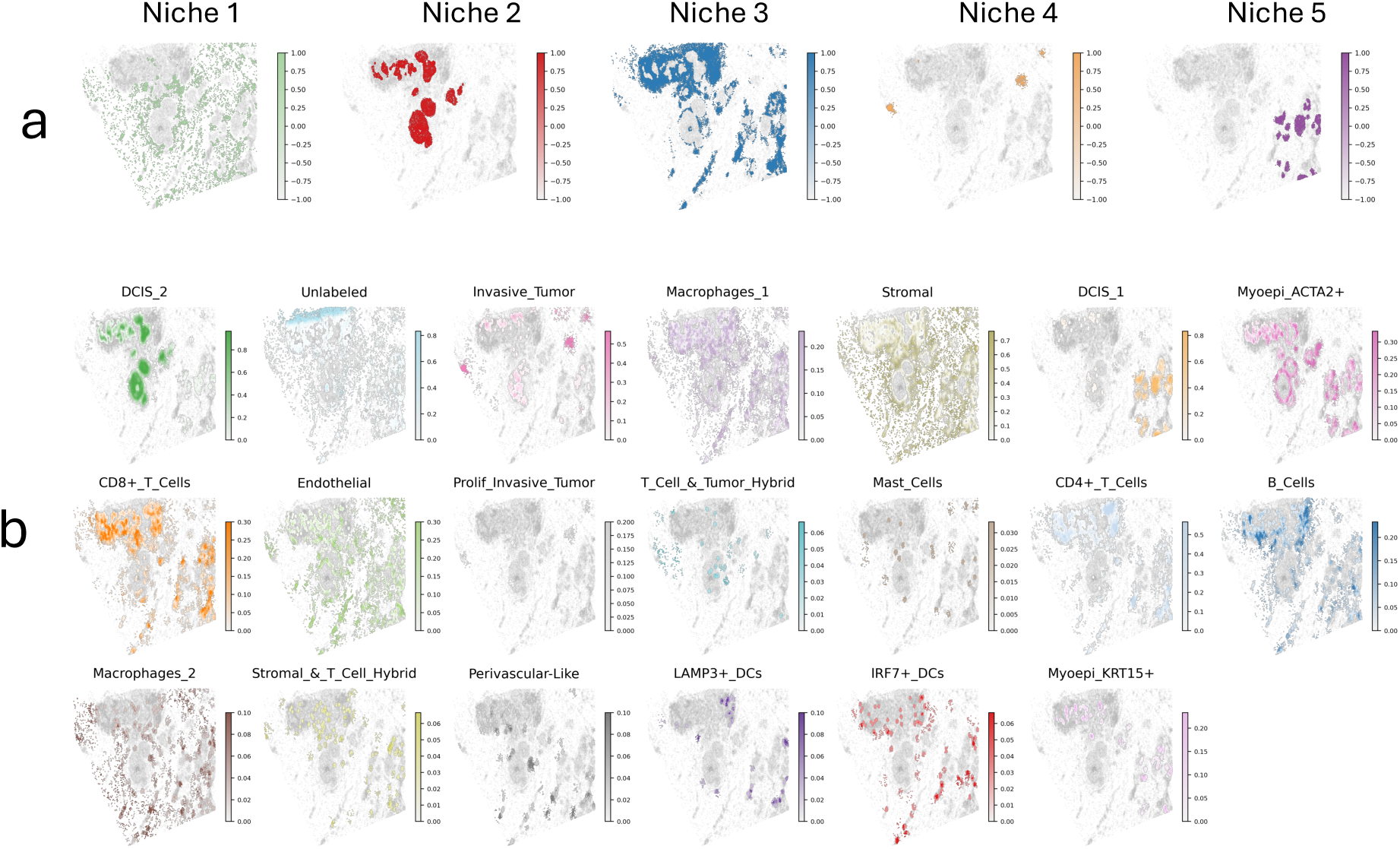
Spatial global (a) and local (b) features used as predictors in S3R, aside from the gene features.

## Notes

### Competing Interest Statement

The authors have declared no competing interest.

### Summary of Updates

Update the latest application and related author information.

